# Re-evaluation of the nodulation capacity of *Sphingomonas sediminicola* DSM 18106^T^ indicates that this strain is not capable of inducing root nodule formation on *Pisum sativum* (pea)

**DOI:** 10.1101/2025.04.30.650786

**Authors:** George C. diCenzo, Samuel M. Gutmanis, Oona Esme, Lionel Moulin

**Author notes:** **Corresponding author:** George C. diCenzo.

## Abstract

Rhizobia are soil-dwelling proteobacteria that can enter into symbiotic nitrogen-fixing relationships with compatible leguminous plants. Taxonomically, rhizobia are divided into alpha-rhizobia, which belong to the class *Alpharoteobacteria*, and beta-rhizobia, which belong to the class *Betaproteobacteria*. To date, all bona fide alpha-rhizobia belong to the order *Hyphomicrobiales*. However, a recent study suggested that *Sphingomonas sediminicola* DSM 18106^T^ is also a rhizobium and is capable of nodulating pea plants (*Pisum sativum*), which would expand the known taxonomic distribution of alpha-rhizobia to include the order *Sphingomonadales*. Here, we attempted to replicate the results of that previous study. Resequencing and computational analysis of the genome of *S. sediminicola* DSM 18106^T^ failed to identify genes encoding proteins involved in legume nodulation or nitrogen fixation. In addition, experimental plant assays indicated that *S. sediminicola* DSM 18106^T^ is unable to nodulate the two cultivars of pea tested in our study, unlike the rhizobium *Rhizobium johnstonii* 3841^T^. Taken together, and in contrast to the previous study, these results suggest that *S. sediminicola* DSM 18106^T^ is not capable of inducing root nodule formation on pea, meaning that the taxonomic distribution of all known alpha-rhizobia remains limited to the class *Hyphomicrobiales*.

## INTRODUCTION

Rhizobia (singular: rhizobium) are soil-dwelling proteobacteria that can enter endosymbiotic relationships with compatible leguminous plants such as pea, common bean, soybean, lentils, alfalfa, and clover. Following an exchange of signals from the plant and rhizobia, the legume forms a specialized organ known as a root nodule within which rhizobia are housed (Gage 2004; Downie 2014; Roy et al. 2020). Generally, the early legume – rhizobium communication involves the production of Nod factors by rhizobia, whose production requires nodulation, or *nod*, genes (D’Haeze and Holsters 2002). However, some rhizobia can induce legume nodulation in a Nod factor-independent fashion; this generally requires the secretion of nodulation outer proteins (known as Nops) by a type III secretion system (T3SS) (Okazaki et al. 2013; Staehelin and Krishnan 2015), although some photosynthetic bradyrhizobia can nodulate plants of *Aeschynomene* genus via an unknown mechanism that is independent of both Nod factors and T3SSs (Giraud et al. 2007; Camuel et al. 2023). Within the nodule, rhizobia convert atmospheric nitrogen gas into ammonia (a process known as symbiotic nitrogen fixation), which is provided to the plant host in exchange for carbon. Under ideal conditions, rhizobia can fulfill the nitrogen demands of the host legume, which has led to rhizobia being commonly applied to legume agricultural fields as a bio-inoculant, largely replacing the requirement for nitrogen fertilizers.

Rhizobia are a polyphyletic group of bacteria distributed between two taxonomic classes: *Alphaproteobacteria* and *Betaproteobacteria* (Wang 2019) (sites.google.com/view/taxonomyagrorhizo/home/species-with-standing-in-nomenclature). Rhizobia within the class *Alphaproteobacteria* are known as alpha-rhizobia, while those within the class *Betaproteobacteria* are known as beta-rhizobia. Sporadic publications have also claimed the existence of rhizobia within the class *Gammaproteobacteria* (Benhizia et al. 2004; Huang et al. 2012), called gamma-rhizobia, but these claims remain controversial as they have not been validated and generally do not fulfill Koch’s postulates. For example, the first study to claim the identification gamma-rhizobia was subsequently disproven by the same group (Benhizia et al. 2004; Muresu et al. 2008). Thus, while it is possible gamma-rhizobia may exist in nature, their existence has yet to be proven.

Within the *Alphaproteobacteria*, all bona fide rhizobia belong to the order *Hyphomicrobiales* (formerly *Rhizobiales*) and are spread across six families (*Rhizobiaceae, Bartonellaceae, Nitrobacteriaceae, Methylobacteriaceae, Xanthobacteriaceae*, and *Devosiaceae*) (Wang 2019; diCenzo et al. 2024). Recently, Mazoyon et al. (2023) proposed that *Sphingomonas sediminicola* DSM 18106^T^ of the family *Sphingomonadaceae* (order *Sphingomonadales*) carries an ∼520 kb plasmid encoding genes required for nodulation and nitrogen fixation, and that it can form nodules on *Pisum sativum* (pea) plants. This result would extend the known taxonomic distribution of alpha-rhizobia beyond the order *Hyphomicrobiales*. However, our own preliminary re-analysis of the *S. sediminicola* DSM 18106^T^ sequencing data, in our roles as members of the Subcommittee on the Taxonomy of Rhizobia and Agrobacteria of the International Committee on Systematics of Prokaryotes, failed to identified nodulation genes (Mousavi and Young 2025), contrary to the results of Mazoyon et al. (2023). In addition, *S. sediminicola* DSM 18106^T^ represents the same original isolate as *S. sediminicola* KACC 15039^T^ (genebank.rda.go.kr/microbeSearchView.do?sFlag=ONE&sStrainsn=30257) but deposited in a different strain collection. Although the genomes of these two strains should therefore be the same, no plasmid is present in the published *S. sediminicola* KACC 15039^T^ genome assembly (NCBI GenBank Assembly accession GCF_014489515.1).

To resolve this situation, we resequenced the genome of *S. sediminicola* DSM 18106^T^ and tested its ability to nodulate two cultivars of pea plants. We report that *S. sediminicola* DSM 18106^T^ does not contain nodulation genes and is unable to form nodules on either cultivar of pea. These results suggest that *S. sediminicola* DSM 18106^T^ is not a rhizobium and are consistent with alpha-rhizobia being taxonomically restricted to the order *Hyphomicrobiales*.

## MATERIALS AND METHODS

### Bacterial strains and growth conditions

*S. sediminicola* DSM 18106^T^ (An et al. 2013) was purchased from the German Collection of Microorganisms and Cell Cultures GmbH (DSMZ) in 2024. *S. sediminicola* DSM 18106^T^ was initially grown using full strength Reasoner’s 2A (R2A) medium (0.5 g/L yeast extract, 0.5 g/L proteose peptone, 0.5 g/L casamino acids, 0.5 g/L glucose, 0.5 g/L starch, 0.3 g/L sodium pyruvate, 0.3 g/L dipotassium phosphate, 0.05 g/L magnesium phosphate heptahydrate, and 15 g/L agar for solid medium) at 28°C. However, we found that *S. sediminicola* DSM 18106^T^ grew very poorly on this medium. Therefore, after the initial recovery of *S. sediminicola* DSM 18106^T^ from the freeze-dried culture, we switched to ½ R2A medium (similar to full strength R2A but with half the amount of all compounds except for the agar), which we found supported much faster growth of this strain.

*Rhizobium johnstonii* 3841^T^ (formerly *Rhizobium leguminosarum* 3841) (Johnston and Beringer 1975; Young et al. 2023) was grown using TY medium (5 g/L tryptone, 2.5 g/L yeast extract, 10 mM calcium chloride, and 15 g/L agar for solid medium) at 28°C.

### 16S rRNA gene sequencing

The 16S rRNA gene of *S. sediminicola* DSM 18106^T^ was amplified using polymerase chain reaction (PCR) with 2X Taq FroggaMix (Froggabio), one colony as a source of DNA, the primers 27f (5’ GAG AGT TTG ATC CTG GCT CAG) and 1495r (5’ CTA CGG CTA CCT TGT TAC GA) (Di Cello et al. 1997). Thirty rounds of PCR were performed with the following cycling conditions: 94°C for 30 seconds, 50°C for 30 seconds, and 72°C for 90 seconds. The amplified PCR product was purified using a Monarch PCR & DNA Cleanup Kit (New England Biolabs) and then sequenced using Oxford Nanopore Technologies (ONT) sequencing at Plasmidsaurus (Louisville, KY, USA).

### Whole genome sequencing

One mL of ½ R2A broth was inoculated using a single colony of *S. sediminicola* DSM 18106^T^ and grown for two days at 28°C. Total genomic DNA was isolated using a DNeasy UltraClean 96 Microbial Kit (Qiagen). A DNA sequencing library was prepared using a Rapid Barcoding Kit 96 V14 (SQK-RBK114.96; ONT) and sequenced on a PromethION R10.4.1 flow cell (ONT) using a ONT P2 Solo sequencer according to the manufacturer’s protocol. Basecalling was performed using dorado version 0.5.1 (github.com/nanoporetech/dorado) with the model dna_r10.3.1_e8.2_400bps_sup@v4.3.0.

### Genome assembly and annotation

Five previously published Illumina read sets (accessions: SRR18460303, SRR18460304, SRR18460305, SRR18460306, SRR18460307) (Mazoyon et al. 2023) were downloaded from the Sequence Read Archive (SRA). Illumina reads were filtered using BBDuk version 39.01 (Bushnell 2014) and trimmed using Trimmomatic version 0.39 (Bolger et al. 2014) with the following parameters: LEADING:3 TRAILING:3 MINLEN:36 SLIDINGWINDOW:4:15. Genome assembly was then performed using Unicycler version 0.5.0 (Wick et al. 2017) with SPAdes version 3.15.5 (Bankevich et al. 2012), for each of the five read sets individually and after combining all five read sets.

For genome assembly using ONT data, the assembly was performed using Flye version 2.9.3-b1797 (Kolmogorov et al. 2019) and polished with medaka version 2.0.1 (github.com/nanoporetech/medaka). The assembly was then checked for contamination using a local copy of the National Center for Biotechnology Information (NCBI) Foreign Contamination Screen (FCS) tool (github.com/ncbi/fcs).

Genome annotation was performed using a local copy of the NCBI Prokaryotic Genome Annotation Pipeline (PGAP) version 2023-10-03.build7061 (Tatusova et al. 2016).

### Genome analyses

The quality of the genome assemblies were checked using CheckM version 1.2.3 (Parks et al. 2015) and taxonomically classified using the Genome Taxonomy Database Toolkit (GTDB-Tk) version 2.4.0 with the R220 database (Chaumeil et al. 2020). The similarity of assembled genomes was measured using FastANI version 1.33. Proteomes were searched for homologs of Nod proteins using BLASTp version 2.9.0+ (Camacho et al. 2009). As query sequences, the amino acid sequences of NodA (accession: Q1M7W9_RHIJ3), NodB (Q1M7W8_RHIJ3), and NodC (Q1M7W7_RHIJ3) were downloaded from UniProt (The UniProt Consortium 2024), while the amino acid sequences of the NodA, NodB, and NodC proteins identified in *S. sediminicola* DSM 18106^T^ by Mazoyon et al. (2023) were copied from Table S3 of their study.

We also used NCBI’s implementation of BLASTn (https://blast.ncbi.nlm.nih.gov/blast) to search the raw Illumina data produced by Mazoyon et al. (2023) (accession numbers SRX14592935 to SRX14592939) for the *nodA, nodB* and *nodC* sequences described in Table S3 of the same article. We used SRX1970119 of *Rhizobium johnstonii* strain VF39 as a positive control for the BLASTn analyses.

### Plant assays

Seeds of two cultivars of *P. sativum* (which are not named on the request of the seed providers) were surface sterilized with 3.5% (v/v) hypochlorite for 20 minutes and then rinsed with sterile ddH_2_O for 60 minutes with the ddH_2_O replaced every 20 minutes. Sterilized seeds were spread on 1X water agar plates, cold stratified at 4°C in the dark for two days, and then left to germinate for three days at 21°C. Seedlings were planted in autoclaved Leonard Assemblies (ref), which consisted of two Magenta Jars connected by a cotton wick that extended from the top jar (containing a 1:1 [w/w] mixture of vermiculite and sand) to the bottom jar (containing 250 mL of Jensen’s medium that was initially poured over the top jar) (Jensen 1942; Leonard 1943). One seedling was planted per Leonard Assembly. The plants were grown in the Queen’s University Phytotron’s greenhouse for two days prior to inoculation with either *S. sediminicola* DSM 18106^T^ (∼2 × 10^6^ colony forming units [CFU]), *R. johnstonii* 3841^T^ (1 × 10^8^ CFU), or with 1mL of ½ R2A or TY media (uninoculated control); these CFU counts were chosen to match the values used by Mazoyon et al. (2023). To prepare the bacterial cells for inoculation, liquid cultures were centrifuged at 16,000 x *g* for one minute, after which the cell pellets were resuspended in 10% of the initial volume of the appropriate growth media (TY or ½ R2A). Cell suspensions were diluted to an optical density at 600 nm (OD600) of 0.75, and one mL added to each pot. Plants were then grown in the Queen’s University Phytotron’s greenhouse for 28 days. At the end of their growth period, plants were visually inspected for the presence of nodules, and then cut to separate the shoots and roots, which were individually dried at 60°C for two weeks prior to measuring their dry weights.

To test if differences in the nodulation capabilities of either *S. sediminicola* DSM 18106^T^ and *R. johnstonii* 3841^T^ were simply due to differences in the inoculum dose, a second trial was run in which cell suspensions were diluted to an OD600 of 2.916 for *S. sediminicola* DSM 18106^T^ and 0.058 for *R. johnstonii* 3841^T^, corresponding to ∼7.77 × 10^6^ CFU for both species. All other methods were the same as the first trial described above.

## RESULTS AND DISCUSSION

### *S. sediminicola* DSM 18106^T^ genomes assembled from published Illumina sequencing reads lack plasmids and nodulation genes

As the *S. sediminicola* DSM 18106^T^ genome sequence assembled by Mazoyon et al. (2023) was not publicly available, we instead downloaded the five raw Illumina read sets made available by the authors. Genome assemblies were then produced using Unicycler with each of the read sets individually or after merging the read sets (**Table 1**). Three of the resulting genome assemblies were classified as *S. sediminicola* by the GTDB-Tk tool. One of the assemblies was classified as *Sphingomonas daechungense*, while the fifth assembly was a mixed assembly of *S. sediminicola* and a *Paenibacillus* species. The combined assembly was classified as a mixed assembly of *S. sediminicola, S. daechungense*, and *Paenibacillus* (**Table 1**). As a result, we focused our downstream analyses on the three assemblies corresponding to pure *S. sediminicola* DSM 18106^T^ cultures. Each of the three assemblies had a total length between ∼2.755 Mb and ∼2.761 Mb (**Table 1**), in line with the genome size of *S. sediminicola* KACC 15039^T^ (∼2.757 Mb; GenBank Assembly accession GCF_014489515.1) and the *S. sediminicola* DSM 18106^T^ chromosome size (∼2.756 Mb) reported by Mazoyon et al. (2023). These results suggest that our re-assemblies of the Illumina read sets generated by Mazoyon et al. (2023) do not contain an ∼ 520 kb plasmid, in contrast to the previous report.

**Table 1.**
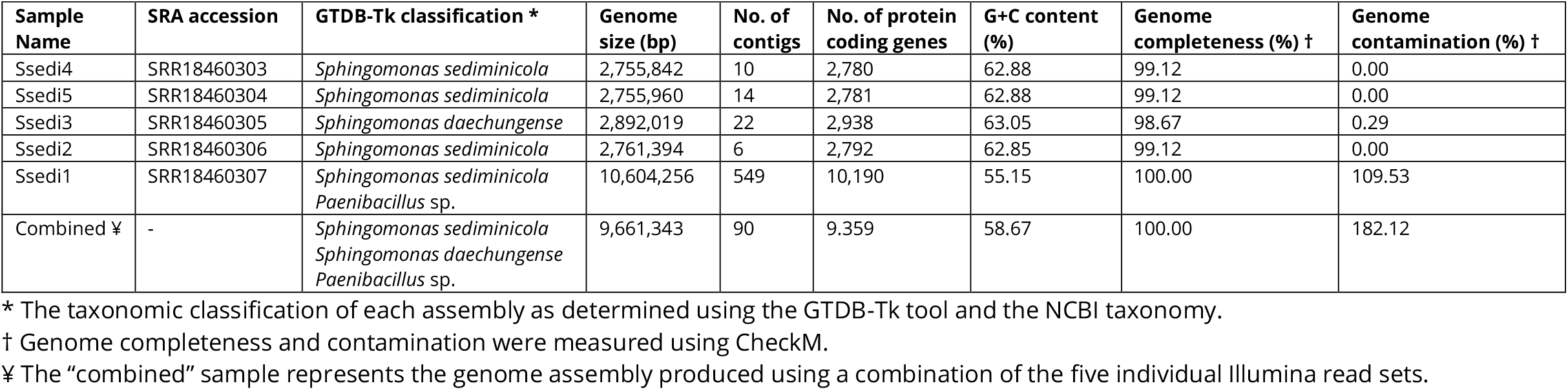
Assembly and annotation statistics of the genome assembled using five published *Sphingomonas sediminicola* DSM 18106^T^ Illumina sequencing read sets.

We next tested whether the proteomes of the three assemblies (generated using the PGAP tool) include proteins involved in symbiotic nitrogen fixation. BLASTp searches of the proteomes failed to identify homologs of the nitrogenase proteins NifH, NifD, and NifK when using as the query the NifH, NifD, or NifK proteins of *R. johnstonii* 3841^T^ or the NifH protein reported as being encoded by *S. sediminicola* DSM 18106^T^ by Mazoyon et al. (2023) (**Dataset S1**); no NifD or NifK were previously reported as being encoded by *S. sediminicola* DSM 18106^T^. Likewise, BLASTp searches failed to identify homologs of the common nodulation proteins NodA, NodB, and NodC regardless of whether we used the NodA, NodB, and NodC proteins of *R. johnstonii* 3841^T^ or those reported as being encoded by *S. sediminicola* DSM 18106^T^ by Mazoyon et al. (2023) (**Dataset S1**). We next wondered whether *S. sediminicola* DSM 18106^T^ might nodulate pea in a Nod factor-independent and T3SS-dependent fashion, and searched for homologs of the SctQ protein that functions as the cytoplasmic ring protein of T3SSs. However, we again failed to identify SctQ homologs in our reassembled *S. sediminicola* DSM 18106^T^ genomes (**Dataset S1**).

In case *S. sediminicola* DSM 18106^T^ does encode a symbiotic plasmid represented in the Illumina data but incorrectly excluded from our genome reassemblies, we also used BLASTn to search the raw Illumina data produced by Mazoyon et al. (2023) for the *nodA, nodB* and *nodC* sequences described in Table S3 of the same article. No hits were obtained, whereas matches were found when searching raw Illumina data produced for *R. johnstonii* VF39 as a positive control. These results suggest that the raw Illumina reads generated by Mazoyon et al. (2023) do not contain reads corresponding to *nodA, nodB*, or *nodC*.

Overall, the results of our re-analysis of the *S. sediminicola* DSM 18106^T^ sequencing data reported by Mazoyon et al. (2023) are inconsistent with *S. sediminicola* DSM 18106^T^ carrying a megaplasmid or encoding the proteins required for symbiotic nitrogen fixation.

### Resequencing of *S. sediminicola* DSM 18106^T^ failed to identify plasmids or nodulation genes

To further explore the genome organization and genome content of *S. sediminicola* DSM 18106^T^, we ordered the strain from the DSMZ strain collection and resequenced its genome from a single colony using ONT sequencing. Initial sequencing of just the 16S rRNA gene confirmed that we received the correct strain from DSMZ. Assembly of the whole genome ONT read set led to the generation of a single circular replicon of 2,756,582 bp, with a genome completeness score of 99.12% and a genome contamination score of 0% as measured using CheckM (**Table 2**). No plasmid was present in the assembly. BLASTp searches of the annotated proteome again failed to identify homologs of the nitrogenase proteins (NifH, NifD, NifK), the common nodulation proteins (NodA, NodB, NodC), or the T3SS protein SctQ, when using as the query the proteins of *R. johnstonii* 3841^T^ or those reported as being encoded by *S. sediminicola* DSM 18106^T^ by Mazoyon et al. (2023) (**Dataset S2**).

**Table 2.**
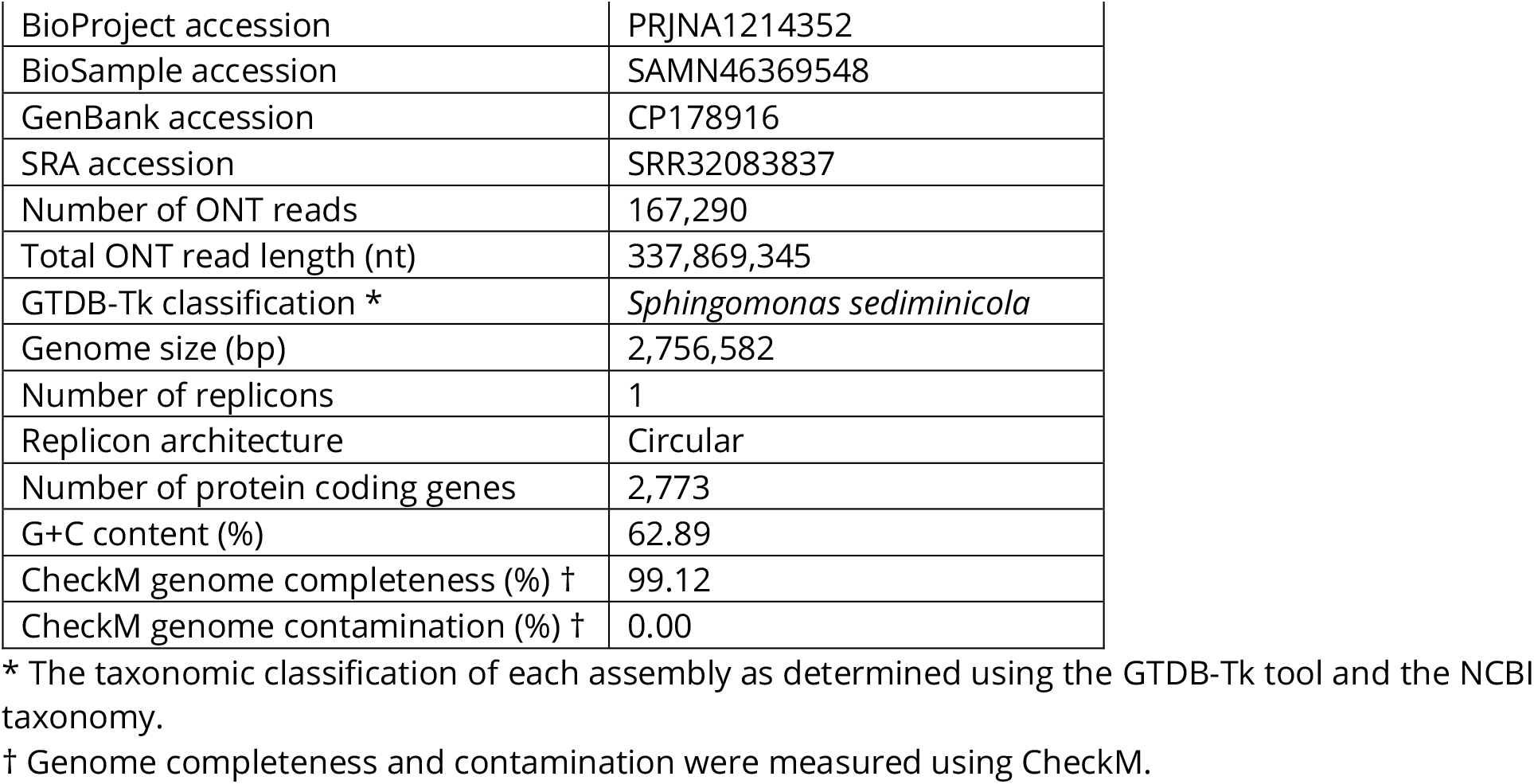
Assembly and annotation statistics of the *Sphingomoans sediminicola* DSM 18106^T^ genome assembled from ONT sequencing data.

Taken together, these results suggest that the *S. sediminicola* DSM 18106^T^ colony we sequenced did not carry a megaplasmid or encode the proteins required for symbiotic nitrogen fixation. This is consistent with the lack of a symbiotic megaplasmid in our re-assemblies of the raw Illumina data generated by Mazoyon et al. (2023), and the lack of a symbiotic megaplasmid in the genome assembly of *S. sediminicola* KACC 15039^T^ (GCF_014489515.1), which represents the same original isolate as *S. sediminicola* DSM 18106^T^ but deposited in a different strain collection. In other words, genome assemblies from three independent whole genome sequencing datasets failed to identify symbiotic megaplasmids, supporting the conclusion that *S. sediminicola* DSM 18106^T^ does not carrying a megaplasmid or encode the proteins required for symbiotic nitrogen fixation.

### *S. sediminicola* DSM 18106^T^ does not induce nodule formation on *P. sativum* (pea)

Lastly, we experimentally tested whether *S. sediminicola* DSM 18106^T^ could induce nodule formation on two cultivars of peas that are grown commercially in Ontario. As a positive control, we inoculated a subset of plants with *R. johnstonii* 3841^T^, a rhizobium known to nodulate pea plants. Four weeks after inoculation, plants inoculated with *R. johnstonii* 3841^T^ were dark green, indicating that the plants received sufficient nitrogen via rhizobial nitrogen fixation (**Figure 1A**). In contrast, plants inoculated with *S. sediminicola* DSM 18106^T^ were comparatively smaller and yellow and looked similar to uninoculated plants (**Figure 1A**), suggesting that little to no nitrogen fixation had occurred. The plant size differences were confirmed by shoot and root dry weight measurements (**Figures 1E, 1F**). Upon uprooting the plants, we observed that whereas plants inoculated with *R. johnstonii* 3841^T^ had large pink nodules (**Figure 1B**), no nodules of any colour were present on the roots of plants inoculated with *S. sediminicola* DSM 18106^T^ (**Figure 1C**) or on uninoculated plants (**Figure 1D**).

**Figure 1.**
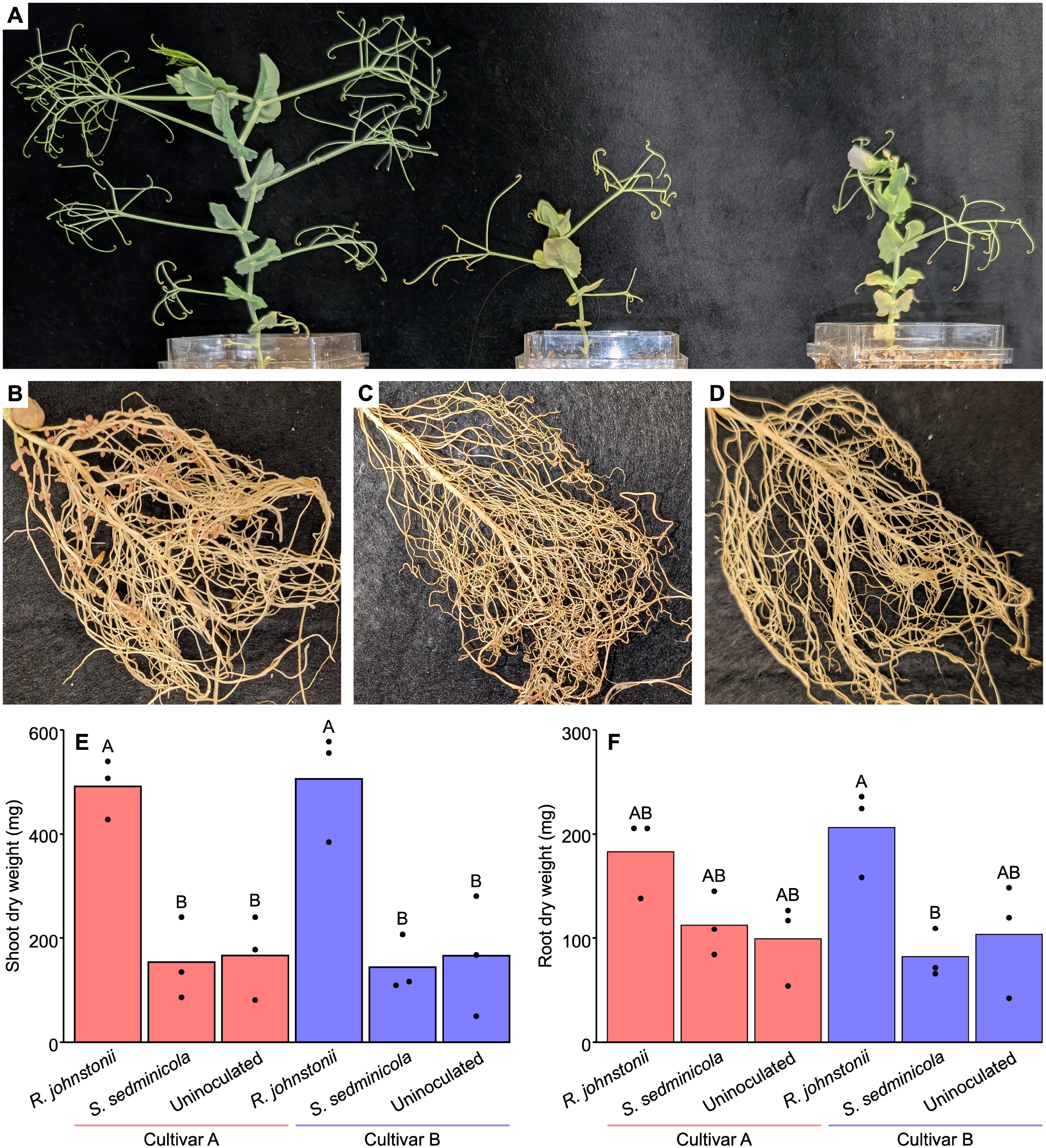
*Sphingomonas sediminicola* DSM 18106^T^ does not nodulate *Pisum sativum* (pea) cultivars. (**A**) Representative photographs are provided of the shoots of *P. sativum* cultivar A plants inoculated with *Rhizobium johnstonii* 3481^T^ (left), *S. sediminicola* DSM 18106^T^ (middle), or no bacteria (right). (**B, C, D**) Representative photographs are provided of the roots of *P. sativum* cultivar A plants inoculated with *R. johnstonii* 3481^T^ (B), *S. sediminicola* DSM 18106^T^ (C), or no bacteria (D). Photographs are not provided for plants of *P. sativum* cultivar B as results are qualitatively similar to those of cultivar A. Graphs showing (**E**) shoot dry weight or (**F**) root dry weight of *P. sativum* plants inoculated with *R. johnstonii* 3841^T^, *S. sediminicola* DSM 18106^T^, or no bacteria. Dots represent individual data points while bars represent the average of the triplicate samples. Statistically significant differences were determined based on a one-way ANOVA followed by a Tukey’s HSD post-hoc test.

Inconsistent with the results of our initial trial, we found numerous nodules on the *S. sediminicola* DSM 18106^T^ inoculated plants upon repeating the experiment, although qualitatively, there tended to be fewer compared to the number of nodules found on *R. johnstonii* 3841^T^ inoculated plants (**Figure S1**). Three nodules per plant were crushed and plated on TY agar (to select for rhizobia) and ½ R2A agar (to allow for growth of *S. sediminicola* DSM 18106^T^). For all 18 nodules, robust growth was observed on the TY agar plates that resembled the morphology of *Rhizobium* spp., whereas little to no growth was observed on the ½ R2A plates. To confirm the identity of the strains isolated on the TY agar plates, the 16S rRNA gene was PCR amplified from six plates (representing the nodule isolates of one nodule per plant) and sequenced. In all cases, the 16S rRNA sequencing data confirmed that the recovered strains belonged to the genus *Rhizobium* and showed 100% identity to the 16S Rrna gene of *R. johnstonii* 3841^T^.

Overall, the results from the two trials suggest that *S. sediminicola* DSM 18106^T^ is unable to nodulate pea plants, and that the nodules formed in the second experiment were due to contamination with a *Rhizobium* strain rather than nodulation by *S. sediminicola* DSM 18106^T^. This is consistent with the genomics data reported above but inconsistent with the experimental results of Mazoyon et al. (2023), who observed the presence of *gusA*-tagged *S. sediminicola* DSM 18106^T^ in pea nodules based on the presence of a blue colour after staining with 5-bromo-4-chloro-3-indolyl glucuronide. As the bacterial nodule population of *S. sediminicola* DSM 18106^T^-inoculated plants was not verified by Mazoyon et al. (2023) using DNA sequencing or by re-isolating bacteria from the nodules, we cannot be certain as to which organism(s) were within the nodules. However, we speculate that in their study, the presence of nodules on *S. sediminicola* DSM 18106^T^-inoculated plants was due to rhizobial contamination with *S. sediminicola* DSM 18106^T^ co-colonizing the nodules as a non-rhizobial endophyte, which is a commonly reported phenomenon (Kosmopoulos et al. 2024; Yu et al. 2025).

## Conclusions

Overall, our results strongly suggest that *S. sediminicola* DSM 18106^T^ is not capable of inducing root nodule formation on pea, contrary to the results of Mazoyon et al. (2023), meaning that the taxonomic distribution of all known alpha-rhizobia remains limited to the class *Hyphomicrobiales*. While we cannot rule out that *S. sediminicola* DSM 18106^T^ can nodulate other pea cultivars, the absence of nodulation genes strongly suggests that it would be unable to do so.

## DATA AVAILABILITY

The raw ONT sequencing reads and the associated annotated genome assembly were deposited at NCBI SRA (accession: SRP559203) and GenBank (accession: CP178916.1), respectively, under the BioProject accession PRJNA1214352, with copies of the files also uploaded to GitHub (github.com/diCenzo-Lab/014_2025_Sphingomonas_sediminicola). Scripts to repeat the genome assembly and annotation, as well as the downstream bioinformatic analyses, are available through GitHub (github.com/diCenzo-Lab/014_2025_Sphingomonas_sediminicola).

## ACKNOWLEDGEMENTS

We thank Rebecca Doyle for kindly providing the pea seeds used in this work, and Turlough Finan for providing *R. johnstonii* 3841^T^. This work was supported by the Natural Sciences and Engineering Research Council of Canada through the Discovery Grants program (RGPIN-2020-0700). This research was also supported by the “Bio-inoculants for the promotion of nutrient use efficiency and crop resiliency in Canadian agriculture” project funded by the Government of Canada through Genome Canada and Genome Prairie (CSAFS-ICT 19308), and the Government of Ontario through an Ontario Research Fund - Interdisciplinary Challenge Teams (ORF-ICT) – Climate Action Genomics Initiative grant (File ICT 19308).

