## Supplementary material for "Re-evaluation of the nodulation capacity of *Sphingomonas sediminicola* DSM 18106^T^ indicates that this strain is not capable of inducing root nodule formation on *Pisum sativum* (pea)": Figure_S1

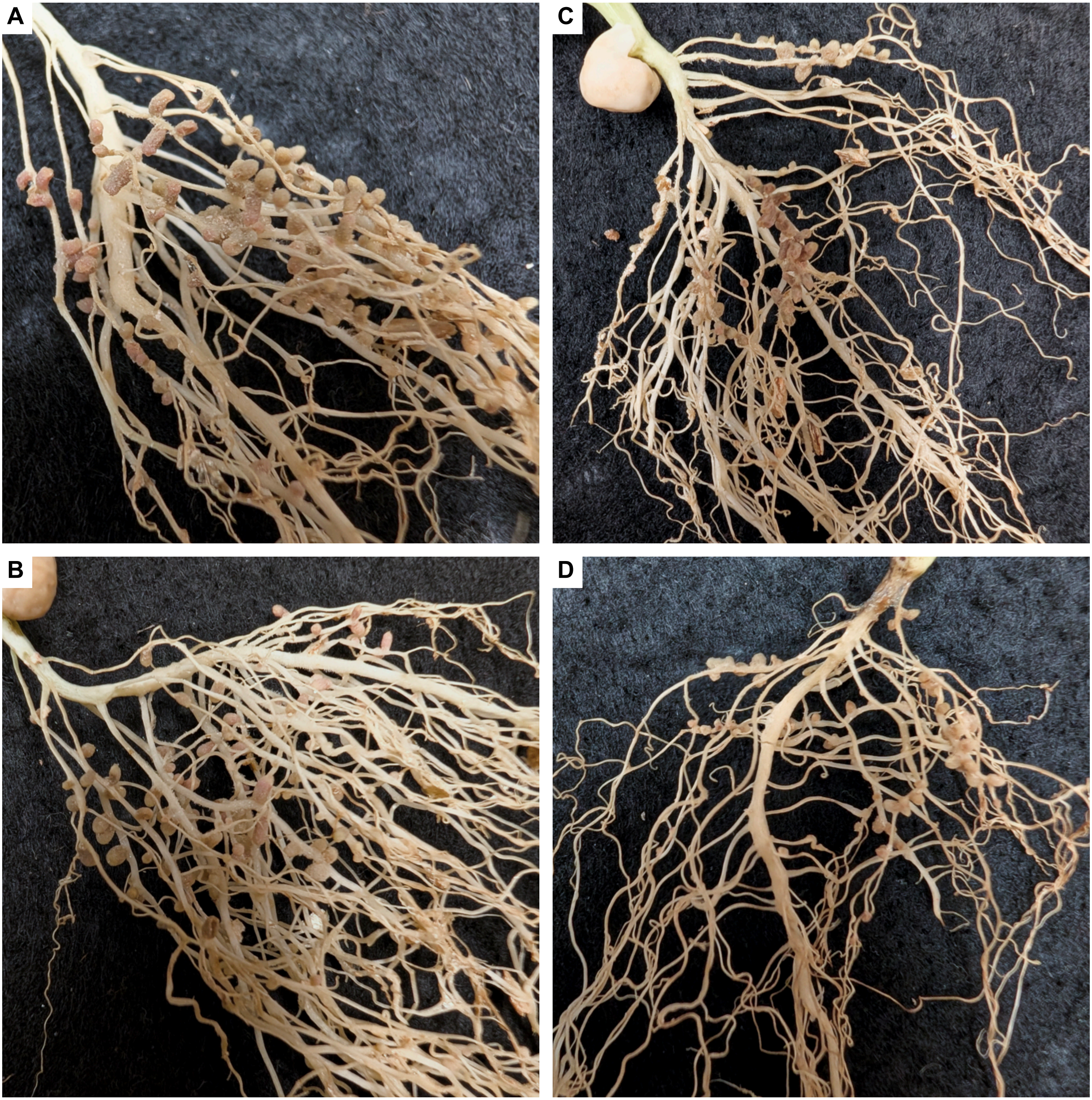


**Figure S1.** Representative photographs of the roots of *Pisum sativum* (pea) cultivar A plants inoculated with (**A**,**B**) *Rhizobium johnstonii* 3841T or (**C**,**D**) *Sphingomonas sediminicola* DSM 18106T. PCR amplification and sequencing of the 16S rRNA gene of six nodule isolates collected from the *S. sediminicola* DSM 18106T inoculated plants indicated that the nodules were colonized by a contaminating *Rhizobium* strain.
